# NON-RANDOM TIMING OF ECOLOGICAL SHIFTS ON CARIBBEAN CORAL REEFS SUGGESTS REGIONAL CAUSES OF CHANGE

**DOI:** 10.1101/672121

**Authors:** William F. Precht, Richard B. Aronson, Toby A. Gardner, Jennifer A. Gill, Julie P. Hawkins, Edwin A. Hernández-Delgado, Walter C. Jaap, Tim R. Mcclanahan, Melanie D. Mcfield, Thaddeus J.T. Murdoch, Maggy M. Nugues, Callum M. Roberts, Christiane K. Schelten, Andrew R. Watkinson, Isabelle M. Côté

**Affiliations:** Marine and Coastal Programs, Dial Cordy and Associates, 1011 Ives Dairy Road, Suite 210, Miami, FL 33179, USA < >; Department of Ocean Engineering and Marine Sciences, Florida Institute of Technology, 150 West University Boulevard, Melbourne, FL 32901, USA < >; Stockholm Environment Institute, Linegatan 87D, Stockholm, Sweden. < >; Centre for Ecology, Evolution and Conservation, School of Biological Sciences, University of East Anglia, Norwich NR4 7TJ, UK < >; Environment Department, University of York, YO1O 5DD, UK < >; Department of Environmental Sciences and Center for Applied Tropical Ecology and Conservation, Applied Marine Ecology Laboratory, University of Puerto Rico, 17 Ave Universidad, STE 1701, San Juan, PR 00925-2537; Sociedad Ambiente Marino, P.O. Box 22158, San Juan, Puerto Rico 00931-2158 < >; Lithophyte Research LLC, 273 Catalan Blvd NE, Saint Petersburg, FL 33704, USA < >; Wildlife Conservation Society, Marine Programs, Bronx, NY 10460, USA < >; Smithsonian Marine Station, Ft Pierce FL 34949 < >; Murdoch Marine Ltd., P.O. Box 513, Warwick, WKBX, Bermuda < >; EPHE, Laboratoire d’Excellence “CORAIL”, PSL Research University, UPVD, CNRS, USR 3278 CRIOBE, F-66360 Perpignan, France < >; GEOMAR, Helmholtz Centre for Ocean Research Kiel, Düsternbrooker Weg 20, 24105, Kiel, Germany < >; Living with Environmental Change, School of Environmental Sciences, University of East Anglia, Norwich NR4 7TJ, UK < >; Department of Biological Sciences, Simon Fraser University, Burnaby BC, V5S 1A6, Canada < >

## Abstract

Caribbean reefs have experienced unprecedented changes in the past four decades. Of great concern is the perceived widespread shift from coral to macroalgal dominance and the question of whether it represents a new, stable equilibrium for coral-reef communities. The primary causes of the shift -- grazing pressure (top-down), nutrient loading (bottom-up) or direct coral mortality (side-in) -- still remain somewhat controversial in the coral reef literature. We have attempted to tease out the relative importance of each of these causes. Four insights emerge from our analysis of an early regional dataset of information on the benthic composition of Caribbean reefs spanning the years 1977–2001. First, although three-quarters of reef sites have experienced coral declines concomitant with macroalgal increases, fewer than 10% of the more than 200 sites studied were dominated by macroalgae in 2001, by even the most conservative definition of dominance. Using relative dominance as the threshold, a total of 49 coral-to-macroalgae shifts were detected. This total represents ∼35% of all sites that were dominated by coral at the start of their monitoring periods. Four shifts (8.2%) occurred because of coral loss with no change in macroalgal cover, 15 (30.6%) occurred because of macroalgal gain without coral loss, and 30 (61.2%) occurred owing to concomitant coral decline and macroalgal increase. Second, the timing of shifts at the regional scale is most consistent with the side-in model of reef degradation, which invokes coral mortality as a precursor to macroalgal takeover, because more shifts occurred after regional coral-mortality events than expected by chance. Third, instantaneous observations taken at the start and end of the time-series for individual sites showed these reefs existed along a continuum of coral and macroalgal cover. The continuous, broadly negative relationship between coral and macroalgal cover suggests that in some cases coral-to-macroalgae phase shifts may be reversed by removing sources of perturbation or restoring critical components such as the herbivorous sea urchin *Diadema antillarum* to the system. The five instances in which macroalgal dominance was reversed corroborate the conclusion that macroalgal dominance is not a stable, alternative community state as has been commonly assumed. Fourth, the fact that the loss in regional coral cover and concomitant changes to the benthic community are related to punctuated, discrete events with known causes (i.e. coral disease and bleaching), lends credence to the hypothesis that coral reefs of the Caribbean have been under assault from climate-change-related maladies since the 1970s.

## Introduction

In their natural state, coral reefs are non-equilibrial, dynamic, disturbance-dominated ecosystems. However, in the past four decades coral reefs around the world have experienced unprecedented ecological changes (Wilkinson 2002, Gardner et al. 2003, Bruno and Selig 2007). Most notable have been a drastic decline in living cover of hard corals and an increase in macroalgal abundance (Aronson and Precht 2001a, 2001b, McManus and Polsenberg 2004). Increases in macroalgae have been interpreted by some authors as the single best indicator of reef degradation (Steneck and Sala 2005, Mumby et al. 2006, 2007).

The first regional-scale, quantitative assessment of the extent of coral loss showed that, in aggregate, Caribbean reefs experienced a decline in coral cover from ∼50% to ∼10%, from 1977 to 2001 (Gardner et al. 2003). Using a larger data set, Schutte et al. (2010) calculated that the average coral cover was around 16% in 2006. More recently, Jackson et al. (2014) noted “that the average coral cover for the wider Caribbean was around 17% in 2012.” On some reefs, the decline of coral cover has been accompanied by an increase in macroalgal cover, leading to a community phase shift *sensu* Done (1992). The prevalence of shifts from the coral-dominated state to a macroalgae-dominated state was reviewed by Bruno et al. (2009). They argued that most reefs around the world are not dominated by macroalgae (with dominance defined as ≥50% cover); rather, most reefs are characterized by macroalgal cover and highly variable coral cover that averages < 50%. Although on many reefs there are more macroalgae today than compared with any recorded historical baseline (see Bruno et al. 2014), the causes of the increase, particularly on Caribbean reefs, are still debated in the literature. Three non-exclusive scenarios have been proposed: the top-down (overfishing), bottom-up (eutrophication and pollution) and side-in (perturbations causing direct coral mortality and habitat loss) models (Precht and Aronson 2006).

The top-down model of reef decline, which suggests that macroalgal dominance resulted primarily from declines in grazing intensity and was derived mainly from observations of Jamaican reefs, holds that phase shifts are primarily a consequence of overfishing (Knowlton 1992, Hughes 1994, Hughes et al. 1999, Jackson et al. 2001, Bellwood et al. 2004). In this model, the depletion of large predators, which occurred centuries ago (Jackson 2001, Jackson et al. 2001); the subsequent switch to overfishing of herbivores (Pauly et al. 1998); and finally the mass mortality in 1983–1984 of the herbivorous sea urchin *Diadema antillarum* (Lessios 1988), the grazing activity of which had concealed the loss of herbivorous fishes (Jackson 2001), triggered a proliferation of macroalgae and concomitant coral mortality (Hughes 1994, Jackson et al. 2001). Although the link between herbivory and macroalgal abundance is clear (Lessios 1988, Aronson and Precht 2000, Williams and Polunin 2001, Idjadi et al. 2010, Adam et al. 2015), the role of macroalgal growth in causing significant coral mortality is ambiguous (Aronson and Precht 2006, Nugues and Bak 2006, see also Alevizon and Porter 2014).

The bottom-up scenario maintains that declining water quality, particularly as a result of nutrient enrichment, causes shifts from coral to macroalgal dominance, as was observed in the classic example from Kaneohe Bay, Hawaii (Banner 1974, Hunter and Evans 1995, Bahr et al. 2015). Lapointe (1997) argued that nutrient enrichment is the main cause of shifts to algae-dominated states on reefs throughout the Caribbean, and others have argued that “the collapse of many Caribbean coral reefs was long preceded by… increased nutrient and sediment runoff from land” (Bellwood et al. 2004). A number of reviews have assessed the role of nutrient enrichment in the decline of corals (Lirman 2001, Szmant 2002, Precht and Miller 2007) and the relationships among nutrients, algae, and algae–coral competition (Hughes et al. 1999, Miller et al. 1999, McClanahan et al. 2004, McManus and Polsenberg 2004, Littler et al. 2006, Nugues and Bak 2006, Furman and Heck 2008, Mumby and Steneck 2008, Sotka and Hay 2009). In the Florida Keys, Lirman and Fong (2007) showed that patterns of coral cover, population size-structure, growth, and mortality are not related to water-quality gradients. These authors have all essentially concluded that the regional losses of coral cover and increases in macroalgal abundance are not due to nutrient enrichment alone but to a variety of causes.

The side-in scenario invokes perturbations such as infectious diseases, outbreaks of predators, bleaching events, tsunamis, earthquakes, hurricane impacts, and hypothermic stress as important causes of ecological change on coral reefs. These types of disturbance are well-documented, direct causes of the loss of coral cover and habitat on Caribbean reefs (Knowlton et al. 1981, Woodley et al. 1981, Stoddart 1985, Aronson and Precht 2001a, 2001b, Gardner et al. 2005; McWilliams et al. 2005, Scheffers et al. 2006, Lirman et al. 2011, Aronson et al. 2012; Toth et al. 2014). Many authors have suggested that coral mortality itself was the crucial precondition for macroalgal dominance (Ostrander et al. 2000, Lirman 2001, McManus and Polsenberg 2004, Aronson and Precht 2001a, 2001b, 2006; Nugues and Bak 2008, Alevizon and Porter 2014). Macroalgae are also perceived as harmful because in high abundance they can reduce coral recruitment (Kuffner et al. 2006, Box and Mumby 2007, Birrell et al. 2008, Ritson-Williams et al. 2009, Idjadi et al. 2010), potentially slowing the recovery of coral populations from natural and anthropogenic disturbances.

Identifying the causes of ecological change on Caribbean reefs is a complex task because they are likely to vary among reefs and include a combination of localized and regional stressors that have acted separately or in concert at different times on different reefs. We explicitly subscribe to the pluralistic view of Quinn and Dunham (1983) that ecology seeks to evaluate the relative importance of the many causes underlying an observed pattern. There is evidence that on some reefs a combination of factors, including top-down, bottom-up, and side-in causes, may act in concert leading to an extreme situation of dominance by macroalgae (Paine et al. 1998, McClanahan et al. 2002, Côté et al. 2013). It should, however, be possible to gain more precise insight into the regional cause(s) of algal dominance by examining the temporal pattern of coral-to-macroalgae phase shifts in relation to large-scale, punctuated disturbances, such as the catastrophic loss of the keystone herbivore *D. antillarum* to a pandemic disease in 1983–1984 and to episodes of coral disease and mass coral bleaching.

In this paper, we quantify the prevalence of coral-to-macroalgae and macroalgae-to-coral shifts across the Caribbean region over three decades. By examining the timing of the phase shifts, we evaluate the relative likelihood of the causes proposed for these transitions. We also comment on the idea that reefs of the Caribbean have become vast carpets of seaweed (Ogden 1995, Jackson et al. 2001, Pandolfi et al. 2005), and we attempt to discern whether coral- and macroalgae-dominated communities are stable endpoints, each resistant to conversion to the other. Clarifying these hypothetical and empirical aspects of reef ecology is critical to developing pragmatic approaches to coral-reef conservation and management.

## Methods

### Data acquisition

Time-series of data on hard-coral and macroalgal cover for reef sites spanning the wider Caribbean basin (Fig. 1) were obtained through electronic and manual literature searches, as well as direct personal communications with reef scientists, site managers and institutional librarians. Electronic literature searches were conducted using the Science Citation Index (SCI) and Aquatic Sciences and Fisheries Abstracts (ASFA) from 1981 to 2001 and 1988 to 2001, respectively, covering a period of extensive changes in benthic composition on coral reefs (Gardner et al. 2003, Côté et al. 2005). All relevant references cited in these publications were also checked. The only selection criterion employed was that the study reported percent cover of either or both target benthic components, with replicated measurements, from a site within the region. Sites were deemed separate if they were defined as such in the study. When a site crossed a steep depth gradient, parts of the replicate transects were pooled into smaller depth-ranges (e.g. Dustan and Halas 1987).

**Figure 1.**
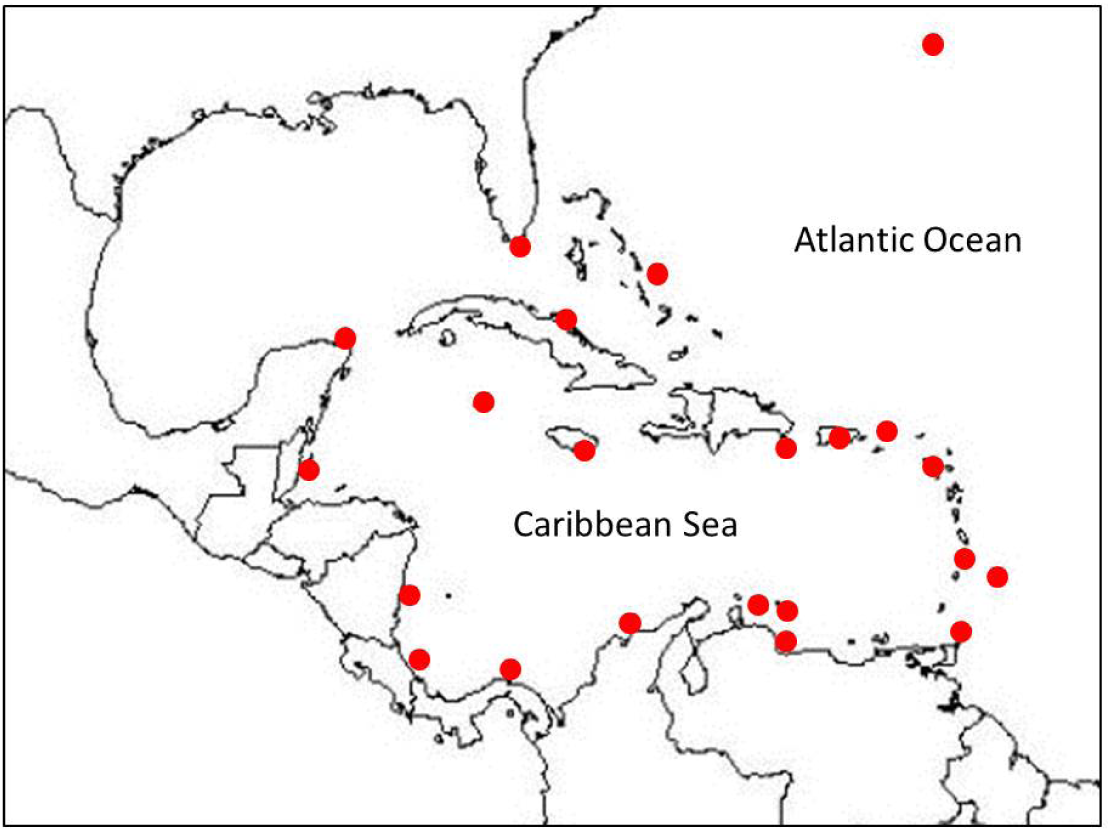
Regional distribution of study sites in the wider Caribbean basin. The study sites from which monitoring data were sourced are shown as solid-red circles. Figure modified from original in Gardner et al. (2003; see Table S1).

### Coral–macroalgae phase shifts and reversals

We defined macroalgae as all erect and anatomically complex algal forms with canopy heights usually > 10 mm (*sensu* Steneck 1988). Crustose coralline algae and algal turfs are also functionally important members of the coral-reef algal assemblage and changes in their relative abundance over time and space may also be critical metrics in diagnosing reef health (e.g. Birrell et al. 2005, Jorissen et al. 2016). However, we did not consider these benthic components in our analysis because many reef-monitoring programs do not differentiate between turfs and other non-erect algal forms (Aronson and Precht 2006, Bruno et al. 2014). Crustose coralline algae, fine algal turfs, and bare space (‘CTB’) are essentially indistinguishable from one another using rapid-assessment monitoring protocols (Aronson et al. 1994, Aronson and Precht 2000).

There is no generally accepted definition of what constitutes a phase shift from corals to macroalgae (Rogers and Miller 2006). The term ‘dominance’ itself is ill-defined in the ecological literature (Bruno et al. 2009, Alevizon and Porter 2014). For example, the definition from Biology Online (2013) is “the species that predominates in an ecological community, particularly when they are most numerous…” But by what comparative threshold do we evaluate the component that is “most numerous?” In quantitative terms, dominance can be a simple majority, which is to say a component (e.g., hard corals or macroalgae) that is more than half of the entire group (i.e., all benthic components combined). By contrast, plurality is the largest subset in a multi-part system. A plurality is not necessarily a majority, as the largest subset may be less than half of the entire group. Finally, relative majority (or relative dominance) is also a subset in a multi-part system but it applies only to one component (e.g., coral) relative to another component (e.g., macroalgae) in the system. At present, there is no consensus in the ecological literature as to whether a simple majority, a plurality, or a relative majority constitutes ecological dominance.

We examined the prevalence of macroalgal dominance in three ways. First, we considered absolute dominance (or majority) of macroalgae. For each year of the dataset, we calculated the proportion of sites with more than 50% macroalgal cover (see also Bruno et al. 2009). Hughes et al. (2010) noted, however, that “dominance” may not be an appropriate metric, because an arbitrary abundance threshold, in this case 50% absolute cover, may be overly conservative. Because macroalgal cover is measured in terms of proportion of the total area rather than the area suitable for colonization (i.e., total area minus the area covered by sand), the usual threshold of 50% may underestimate dominance. We therefore also calculated the proportion of sites with more than 40% macroalgal cover, which would be equivalent to 50% cover where only 80% of the total area surveyed was available for colonization (i.e., average sand cover was 20%, a figure at the high end of our collective personal experience of benthic surveys).

Second, we examined relative dominance (or relative majority) by calculating the proportion of sites in each year with more macroalgae than coral. Both of these methods estimate the prevalence of macroalgal dominance, but they have four limitations. (1) They do not account for the fact that sites may have lost coral cover and/or gained macroalgal cover without having become dominated (in either absolute or relative terms) by the latter while they were monitored. (2) They do not effectively capture changes occurring at sites with naturally low coral cover and high macroalgal cover (see Vroom et al. [2006] for a Pacific example), which may have been that way historically due to regular perturbation. (3) They also do not capture seasonal or local variations in macroalgal cover (Lirman and Biber 2000). (4) Finally, they do not account for other classes of benthic organisms (octocorals, sponges, zoanthids, turfs, crustose coralline algae, etc.) that may have higher absolute cover than either corals or macroalgae or that may ascend to dominance on some temporal scale (e.g., Aronson et al. 2012, Ruzicka et al. 2013, Toth et al. 2014, Tsounis and Edmunds 2017).

Third, due to a paucity of data, we are unsure what the natural cover of coral and macroalgae was on undisturbed Caribbean coral reefs prior to the 1970s (Bruno et al. 2014, Eddy et al. 2018). Knowing this baseline metric would be helpful in interpreting change on reefs through time and developing quantitative targets for management and restoration. Some authors, however, have argued that coral and macroalgal cover by themselves are not reliable metrics of reef health and resilience and that cover only becomes a definitive indicator of phase shifts if the same site is monitored for multiple years (Hughes et al. 2010). To overcome this possible limitation, we examined site-specific ratios of coral to macroalgae at the start and end of each time-series to generate a more dynamic metric of dominance. We recorded the proportion of sites that had moved towards having proportionally less coral than macroalgae, as well as the specific year in which any change of relative dominance was observed at a site. We counted the number of sites in each year that had two attributes: (1) more coral than macroalgae (so they could have shifted); and (2) data for the previous year (so a shift could have been detected). We expressed that number of sites for each year as a proportion of all such sites, and then multiplied by the number of observed shifts (which was 49). This procedure distributed the shifts in proportion to the availability of sites that could shift.

The year of shift could only be identified at sites for which we had concurrent yearly estimates of both coral and macroalgal cover that were separated by no more than one year. Of the 208 sites in the database, 193 met this criterion. To evaluate whether more shifts were observed in each year than expected by chance alone, we used *G*-tests to compare the number of shifts recorded in any given year to the number expected given the total number of shifts observed across all years and the number of sites available to assess a shift in that year (expressed as the proportion of all sites across all years for which we had both coral and macroalgal cover data in that year and the previous year). To account for multiple tests, we adjusted the critical α using the simple Bonferroni correction. The adjusted significance threshold was α_adj_ = 0.002. Finally, we recorded ‘phase-shift reversals,’ when a site that had exhibited greater cover of macroalgae than of coral, either at the start of the study or for at least two consecutive years during the study, shifted to having a higher cover of coral than of macroalgae at the end of its monitoring period.

## Results

A total of 193 reef sites from 71 separate studies across the Caribbean reported concurrent cover of hard corals and macroalgae. The sites were distributed evenly around the Greater Caribbean (similar to the distribution shown in Gardner et al. [2003], with the addition of sites in St Lucia and Saba; Fig. 1). Overall, the time-series of coral and macroalgal cover spanned the years 1977 to 2001, although individual time-series were on average 6 (± 4.7 SD) years long.

When coral-to-macroalgae phase shifts are defined by absolute (or majority) dominance—that is, shifts to ≥50% cover of macroalgae—the prevalence of phase shifts appears to have varied since 1977. Prior to 1981, all sites that were surveyed at the time (*n* = 19) had less than 50% cover of macroalgae. The proportion of sites dominated by macroalgae peaked in 1987 and then decreased to fewer than 10% of sites since 1994 (Fig. 2). This change through time is related in part to the diminishing proportion in the dataset of Jamaican sites, which experienced a unique series of disturbances and ensuing ecological changes in the early 1980s (Côté et al. 2013). Setting the threshold for absolute dominance of macroalgae to 40% cover does not change the temporal pattern or proportion of sites dominated by macroalgal cover.

**Figure 2.**
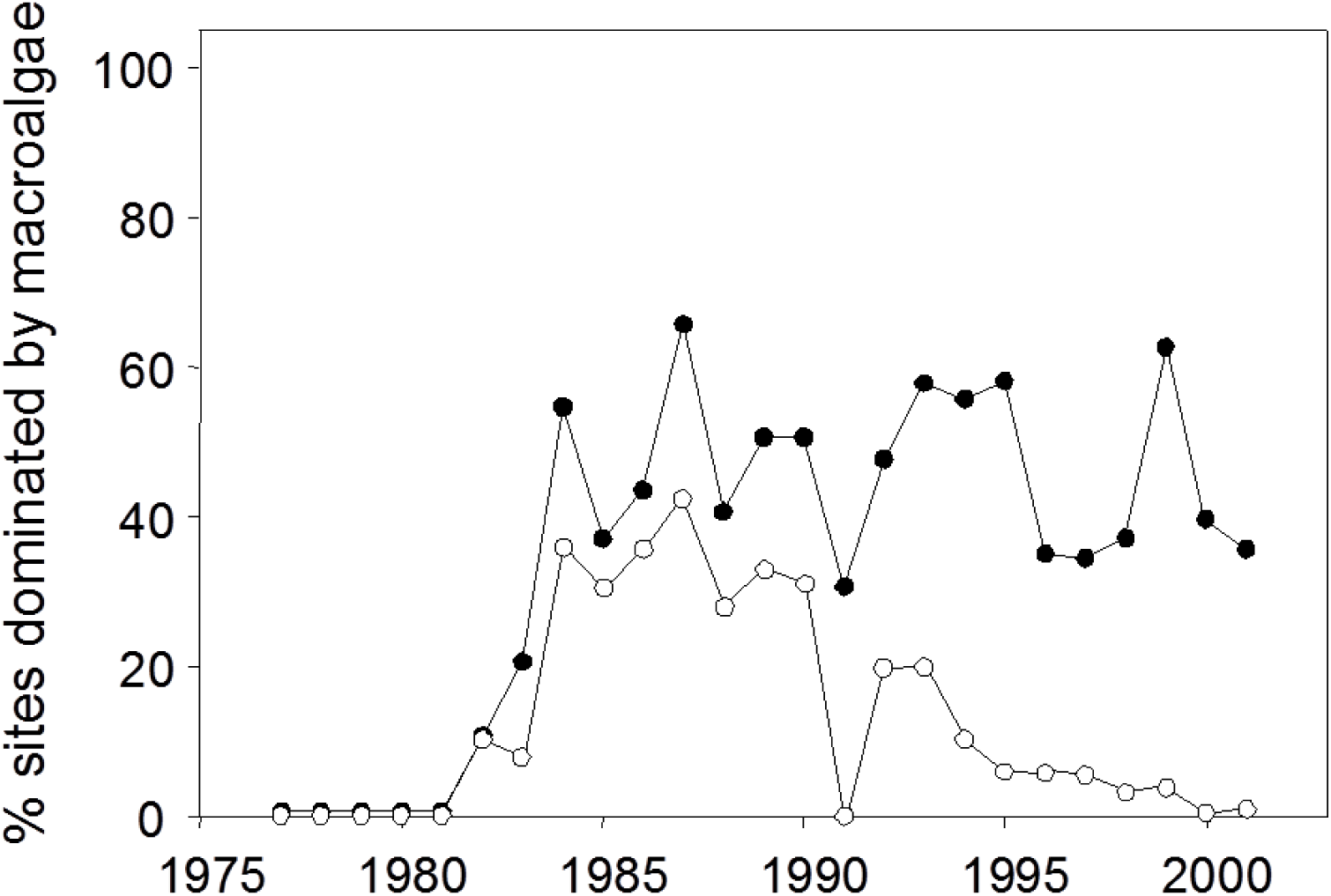
Prevalence of macroalgal dominance in Caribbean reef sites in each year between 1977 and 2001. Macroalgal dominance is defined either in absolute terms (> 50% cover; open circles) or relative terms (higher macroalgal than coral cover; solid circles).

The prevalence of phase shifts is higher if phase shifts are defined as occurring when macroalgal cover exceeds coral cover (i.e., relative dominance of macroalgae). In the late 1970s, all sites exhibited higher cover of hard corals than of macroalgae (Fig. 2). Using this more liberal definition, by 1987 approximately half the sites had more macroalgae than coral, a proportion that has since remained relatively constant. The site-specific analysis reveals an even greater prevalence of phase shifts. Overall, 72% of individual sites moved towards proportionally less coral than macroalgae during their monitoring periods. The difference between coral and macroalgal cover declined until the early 1990s and has remained relatively stable since then at the majority of sites (Fig. 2; see similar analysis in Jackson et al. 2014).

A total of 49 coral-to-macroalgae shifts in relative dominance were detected. This total represents ∼35% of all sites that were dominated by coral at the start of their monitoring periods. Four shifts (8.2%) occurred because of coral loss with no change in macroalgal cover, 15 (30.6%) occurred because of macroalgal gain without coral loss, and 30 (61.2%) occurred owing to concomitant coral decline and macroalgal increase. The shifts occurred in 14 of the 25 years of the study period (Fig. 3). The years posting significantly more shifts to macroalgae than expected by chance were 1983, 1984, 1990 and 1998 (Fig. 3). Note that adjustment of α to maintain a familywise error rate of 0.05 yielded the same results using the conservative simple-Bonferroni procedure and the less conservative sequential-Bonferroni and false-detection-rate procedures (Benjamini and Hochberg 1995). In 1983 and 1984, two-thirds of shifts were caused by macroalgal increase with no change in coral cover, whereas in the 1990 and 1998, 81% of shifts involved coral loss, either with or without macroalgal increases (*G* = 4.62, df = 1, *P* = 0.03).

**Figure 3.**
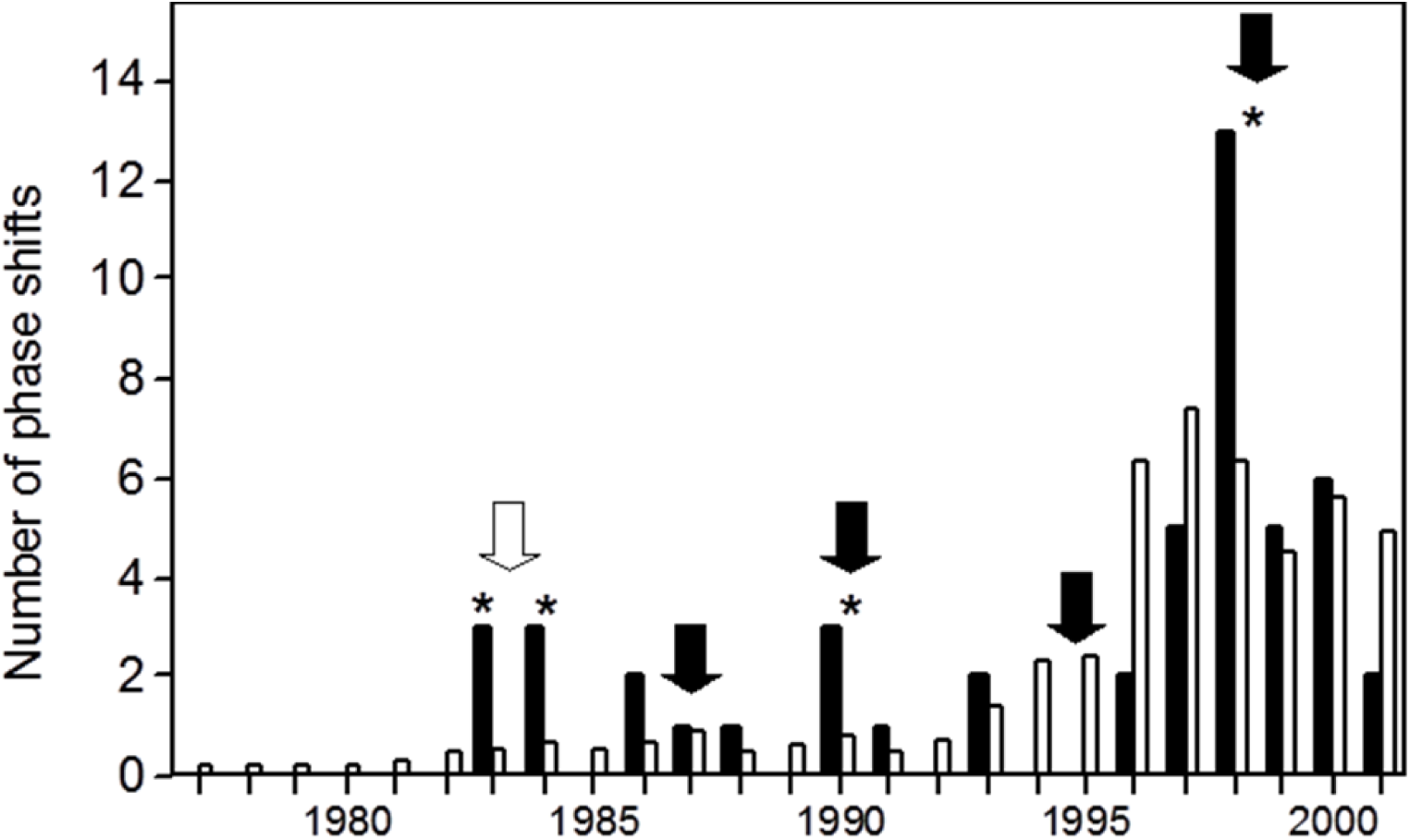
Numbers of coral-to-macroalgae shifts, expressed in relative terms, on Caribbean coral reefs (black bars) each year from 1977 to 2001. Also shown (white bars) are numbers of coral-to-macroalgae shifts expected to occur if these shifts simply occurred in proportion to the number of sites available to shift towards macroalgae dominance each year in the database. * indicates a significantly greater number of observed shifts (G-tests, Bonferroni-corrected α = 0.002). The white arrow denotes the year of mass mortality of the sea urchin *Diadema antillarum*. The black arrows indicate years of mass coral bleaching, the most severe of which was 1998.

Of 57 sites that had more macroalgae than coral at the start of their monitoring periods, 52 (91%) had more macroalgae for the duration of monitoring; however, five reefs showed a clear reversal of the usual coral-to-macroalgae sequence of relative dominance. These five reefs represent ∼5% of all reefs that could have shifted to coral dominance: the 57 sites that were initially macroalgae-dominated, plus 49 sites that had shifted to more macroalgae during the time they were monitored. The reversals all occurred in the 1990s (1994, 1995, 1998 [2 sites], and 1999) and were geographically widespread (Mexico, Colombia, Saba [2 sites], and Puerto Rico). In three cases, the reversals were the result of decreased macroalgal cover only, whereas in two cases, macroalgae decreased and corals simultaneously increased.

## Discussion

Coral-reef degradation and phase shifts are highly variable processes that involve different combinations of coral loss and macroalgal gain in time and space (Bruno et al. 2009). Although we found that nearly three-quarters of reef sites across the Caribbean region changed in the general direction of having less coral and more macroalgae, our analysis also reveals that most Caribbean reefs were not overrun by macroalgae. After 1995, fewer than 10% of sites were dominated by macroalgae in absolute-majority terms. In this respect, our results are remarkably similar to the global analysis of Bruno et al. (2009), who came to a similar conclusion about the extent of macroalgal dominance in the Caribbean using different data.

The timing of shifts from coral to macroalgae sheds light on the causes of these ecological changes. The top-down model of reef decline, which suggests that macroalgal dominance resulted primarily from declines in grazing intensity, implies that the mass mortality of *Diadema antillarum* in 1983–1984 should have had a detectable effect on the timeline of coral–macroalgae shifts in the Caribbean. In 1983, a water-borne pathogen decimated populations of this important herbivore, causing the loss of ∼93% of the *Diadema* throughout the region in less than one year (Lessios 1988, 2016). The mass mortality triggered cascading effects on the reef benthos, as well as on herbivorous and predatory fishes (Hughes et al. 1987, Robertson 1991). We found that in 1983 and 1984 more reefs exhibited shifts towards macroalgae than expected by chance alone, suggesting a regional link between decreased herbivory and increased macroalgal cover. During this two-year period, however, observed phase shifts were caused mainly by increases in macroalgae and not decreases in coral cover. This result calls into question the claim of Jackson et al. (2001) that the *Diadema* pandemic was “the principal cause of coral mortality” throughout the Caribbean.

The early years of our time-series are heavily influenced by Jamaica. The independent contributions of reduced herbivory and coral mortality are, unfortunately, difficult to partition because other events immediately preceded and overlapped the loss of *Diadema* in Jamaica. Most notable were Hurricane Allen in 1980, which caused extensive coral mortality (Woodley et al. 1981), and subsequent losses of coral due to predation and white-band disease (Knowlton et al. 1981). Even in Jamaica, there appears to have been an interaction between coral mortality and reduced herbivory in the shift from coral to macroalgal dominance (Hughes 1994, Andres and Witman 1995, Precht and Aronson 2006). Jamaican reefs had much more coral than macroalgae until 1980, after which the relative difference in cover between these two benthic components began to decline markedly. By 1984, macroalgal cover exceeded coral cover on Jamaican reefs. The difference between macroalgal and coral cover peaked in 1992, with reefs having had on average 70% higher absolute cover of macroalgae than living coral. Corals recovered and macroalgae declined between 1992 and 1999, the last year of our dataset for Jamaica (Fig. 4). The pattern for other Caribbean reefs was superficially similar but quantitatively different. On non-Jamaican reefs, the difference between coral and macroalgae also declined from 1984 to 1994, but less-so, and remained relatively stable for the remainder of the time-series (Fig. 4).

**Fig. 4.**
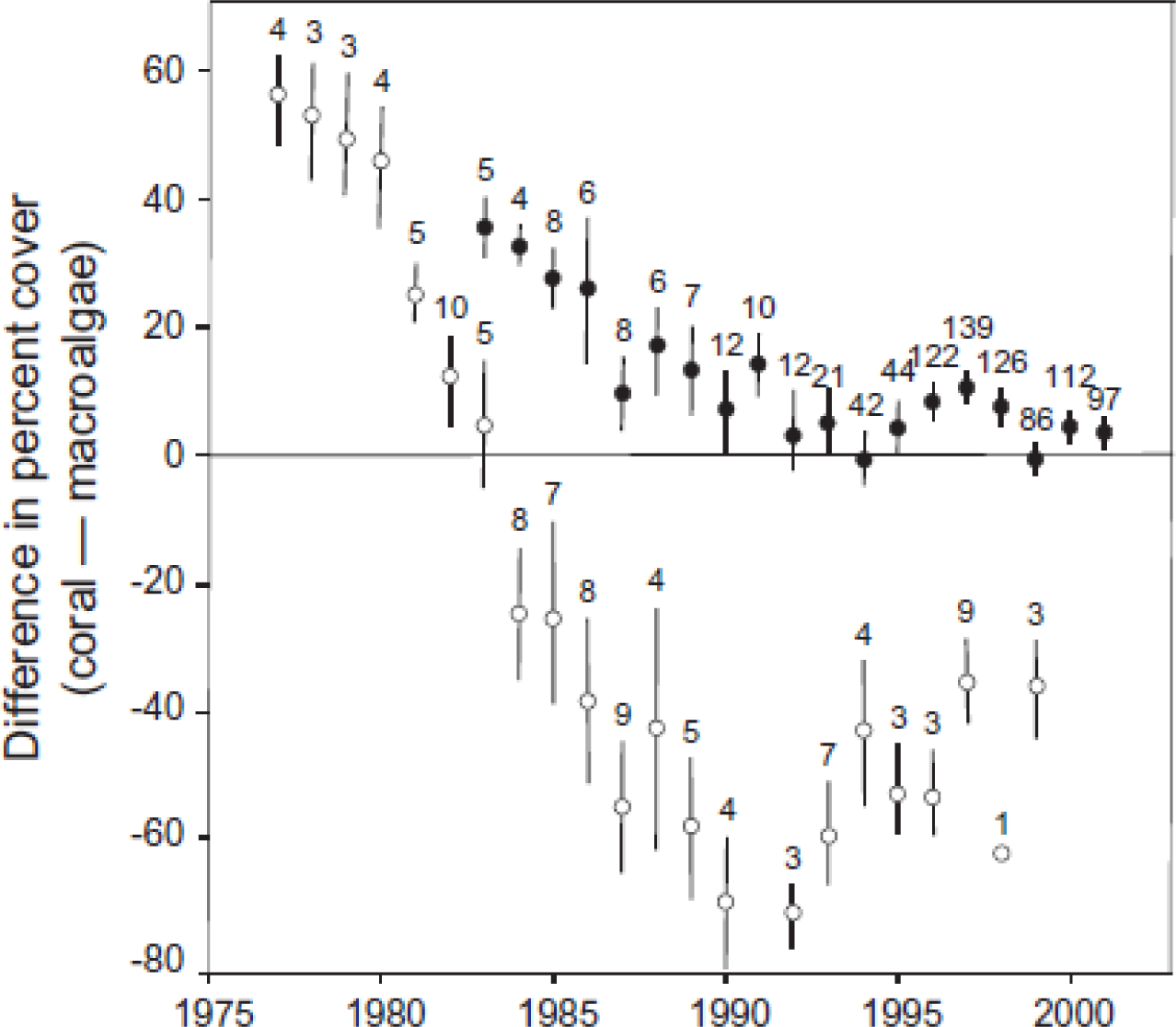
Magnitudes of the difference between percent cover of live coral and macroalgae on Caribbean reefs in each year between 1977 and 2001. Means (±1 SE) are shown separately for Jamaican sites (open circles) and all other sites (filled circles). Positive values indicate that the cover of coral was higher than that of macroalgae. Numbers above the error bars show the number of sites contributing to each mean. Figure modified from original in Côté et al. (2014).

At the regional scale, as in Jamaica significant losses of acroporid corals from white-band disease preceded and overlapped the loss of *Diadema* (Gladfelter 1982, Aronson and Precht 2001a, 2006, 2016). The years 1977–1983 showed significant losses of coral, with up to 40% decline in coral cover (Gardner et al. 2003). The timing of coral loss is consistent with white-band disease having been the primary cause of coral mortality in the Caribbean (Gardner et al. 2003, Aronson and Precht 2006, Alvarez-Filip et al. 2009, Schutte et al. 2010, Aronson and Precht 2016, Kuffner and Toth 2016). The loss of coral cover *per se* during this period was not coincident with widespread coral-to-macroalgae phase shifts, highlighting the importance of *Diadema* in ameliorating the impacts of macroalgae in the face of large-scale disturbances. Once *Diadema* were removed from the system in 1983–1984, however, much of the space made available by catastrophic mortality of the acroporids was rapidly colonized by macroalgae. The two other years, 1990 and 1998, in which there were significantly more coral-to-macroalgae shifts than expected—virtually all of which involved declines in coral cover—were years of mass coral-bleaching events in the Caribbean (McWilliams et al. 2005, Gill et al. 2006). Coral mortality was followed by macroalgal overgrowth of the dead-coral surfaces. The major coral-bleaching event in 1997–1998 resulted in widespread loss of coral, not only in the Caribbean (Aronson et al. 2002a) but also in other regions of the world (Lough 2000, Wilkinson 2002). The trend in coral cover followed a step-type function driven by coral disease and bleaching events as the proximal drivers of coral mortality (Aronson and Precht 2006). The ecological changes occurring in these years are, therefore, most consistent with the side-in model of reef degradation, which invokes coral mortality as the primary factor initiating phase shifts (Precht and Aronson 2006). Mass coral-bleaching events in 1987 and 1995 resulted in only localized areas of coral mortality (Lang et al. 1992, McField 1999, Ostrander et al. 2000). The latter two events were undetectable at the regional scale in our analysis (Fig. 3).

Our results suggest a limited role for changes in grazing pressure in driving phase shifts after 1990. Although we do not have site-specific estimates of herbivory, a regional analysis of Caribbean reef-fish abundance showed declines in herbivorous-fish density over the duration of our study (Paddack et al. 2009). Since 1984 there has been no significant regional-scale recovery of *Diadema* (Hughes et al. 2010, Levitan et al. 2014, Rogers and Lorenzen 2016), suggesting that the intensity of herbivory has been declining on most Caribbean reefs. The temporal patterns in the prevalence of phase shifts—roughly stable or declining since the mid-1980s, depending on the definition of dominance (Fig. 2)—are not consistent with a regional reduction in herbivory. It is possible that reduced herbivory, caused by historical overfishing, set the stage for the extreme impact of coral mortality, as suggested by Jackson (2001, 2014). In a similar vein, Hughes et al. (2012) argued that the regime shifts observed across the Caribbean unfolded so slowly and gradually as to be imperceptible until a tipping point was reached. However, the non-random timing of phase-shifts across a broad gradient of fishing pressure and human influences (Kramer 2003, Newman et al. 2006), and the combined effects of two lethal, essentially concurrent epizootics with regional consequences unprecedented in at least the last several thousand years (Sheppard 1993, Knowlton 2001, Aronson et al. 2002b), argue against such undetected dynamics.

Coral-to-macroalgae phase shifts on Caribbean reefs have been postulated to represent transitions from one alternative stable state to another (Knowlton 1992). Whether the coral- and macroalgae-dominated states are stable alternatives, each of which resists conversion to the other, or whether the coral-to-macroalgae transition is a reversible phase shift (Petraitis and Dudgeon 2004) is critical to determining the resilience of reef communities in the Caribbean (Scheffer and Carpenter 2003, Bellwood et al. 2004). Degraded reefs initially showed no evidence of recovery from a macroalgae-dominated state (e.g., Rogers and Miller 2006). Later literature, however, described a number of phase-shift reversals in the Caribbean including at highly degraded sites in Jamaica (Idjadi et al. 2006). We found evidence of shifts away from macroalgal dominance at five sites during the 1990s, where large changes in herbivory by fish were not reported. Since 2001, reduced macroalgal cover and then increased recruitment of juvenile corals, both associated with the local reappearance of *Diadema*, have occurred, first on Jamaican reefs (Edmunds and Carpenter 2001, Cho and Woodley 2002, Bechtel et al. 2006, Precht and Aronson 2006, Idjadi et al. 2010) and then on reefs elsewhere across the Caribbean (Macintyre et al. 2005, Carpenter & Edmunds 2006, Myhre and Acevedo-Gutiérrez 2007). A full reversal to the coral-dominated state was noted on one Jamaican reef (Idjadi et al. 2006, Precht and Aronson 2006, Crabbe 2009), where *Diadema* had recovered to half their pre-mortality densities (Edmunds and Carpenter 2001) without noticeable changes in fish populations (Woodley and Sary 2002, Hardt 2008) or improvements in water quality (Cho and Woodley 2002, Webber et al. 2005, Greenaway and Gordon-Smith 2006).

There is little evidence to support the notion of discrete, low coral/high macroalgae and high coral/low macroalgae alternative states; in our study, the abundances of hard corals and macroalgae exhibited a continuous, broadly negative relationship (see also Lowe et al. 2011). Reefs with little coral had highly variable amounts of macroalgae, just as reefs with little macroalgae varied greatly in coral cover (Fig. 5). It was more common for Caribbean reefs to shift from coral to macroalgal dominance, with 35% of sites that could do so having done so during the study period, than to shift in the reverse direction, with only 5% of eligible sites having done so. This asymmetry in the direction of shifts is more parsimoniously explained by (1) the relatively short time-series of our data set, (2) the catastrophic and escalating nature of recent disturbances, many of which have caused (and continue to cause) extensive coral mortality (Miller et al. 2009, Eakin et al. 2010, Precht et al. 2016), (3) the rapid colonization and turnover times of the replacement algal species (McCook et al. 2001), and (4) the slow natural succession of long-lived reef fauna following regional, catastrophic mortality of the major foundation species (*Acropora palmata* and *A. cervicornis*) and the keystone herbivore (*Diadema antillarum*)—than by resistance to recovery from the macroalgae-dominated state (Dudgeon et al. 2010). Clearly, if coral-mortality events become more frequent and more severe, the likelihood of complete recovery between disturbance events will diminish (see Palumbi et al. 2008).

**Figure 5.**
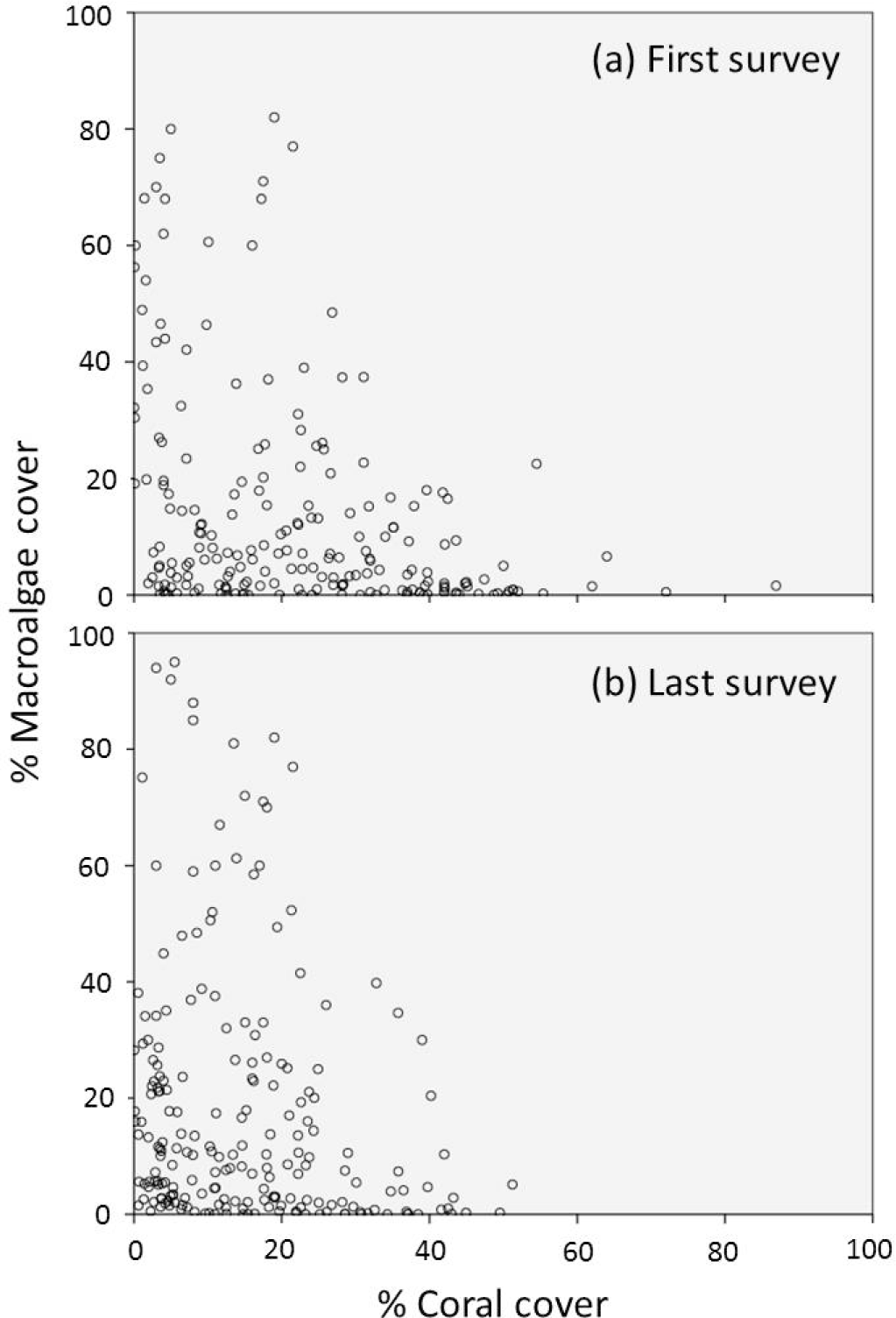
Relationship between macroalgal and coral cover on the reefs included in the study. Data are from the first (a) and last (b) survey of each reef only. The relationship between macroalgal and coral cover is virtually identical. Note the absence of two distinct clusters of reefs, one with high coral and low macroalgal cover and the other with low coral and high macroalgal cover, which would have been predicted by the hypothesis of alternative stable states.

We anticipate that if *Diadema* populations continue to increase across the region, so will the number of phase-shift reversals (Precht and Precht 2016; but see Levitan et al. 2014). Early evaluations suggest, however, that reversing phase shifts may depend on successfully reducing the causative stressors (Dudgeon et al. 2010) and having sufficient rates of activity by functional groups that play key roles in reversing the changes (McClanahan et al. 2011b, Francis et al. 2019). Given enough time and in the absence of additional, significant coral mortality, recovery of Caribbean coral populations seems plausible. Devising restoration and management programs that enhance or restore critical components of the ecosystem, reduce macroalgae, and promote the success of corals should foster resilience and accelerate recovery, at least locally (Mumby 2006, Newman et al. 2006, Precht and Aronson 2006, Hoegh-Guldberg et al. 2007, Furman and Heck 2009, Idjadi et al. 2010, Smith et al. 2010, Edwards et al. 2011, Sandin and McNamara 2012, Jackson 2014, Adam et al. 2014).

Evidence suggests, however, that overfishing is not the major threat to benthic assemblages on coral reefs, nor is the recovery of parrotfish the solution (Aronson and Precht 2006, Rotjan et al. 2006, Russ et al. 2015, Kuffner and Toth 2016, Precht and Precht 2016, Suchley et al. 2016, Bruno et al. 2019). For example, reefs in the Florida Keys, which have maintained relatively high populations of herbivorous fishes (Bohnsack et al. 1994, Alevizon and Porter 2014), should have exhibited considerably less coral loss than reported for the rest of the Caribbean region. Coral loss in Florida, however, was significantly greater than the Caribbean average throughout this period (Schutte et al. 2010), while coral recruit survival and reef recovery were limited (Toth et al. 2014, van Woesik et al. 2014). Protecting fish stocks does not necessarily reduce the cover of macroalgae, increase coral populations, or preserve or increase the topographic complexity that is critical to maintaining and increasing those fish stocks (McClanahan et al. 2011a, 2011b, Bood 2006, Idjadi et al. 2006, Coelho and Manfrino 2007, Kramer and Heck 2007, Ledlie et al. 2007, Mora 2008, Alvarez-Filip et al. 2009, Myers and Ambrose 2009, Stockwell et al. 2009, Selig and Bruno 2010, Alvarez-Filip et al. 2011, Huntington et al. 2011, Lowe et al. 2011, Żychaluk et al. 2012, Reyes-Bonilla et al. 2014, Toth et al. 2014, Cox et al. 2017, Bates et al. 2019).

## Summary

The timing of macroalgal phase-shifts at the regional scale spanning the years 1977– 2001 is most consistent with side-in model of reef degradation, which invokes coral mortality as a precursor to macroalgal takeover, because more shifts occurred after regional coral-mortality events than expected by chance. The implication of this conclusion is that regional causes of coral mortality far outweighed local causes. Why is this important? Reversing coral decline will depend on simultaneously addressing the proximal causes of regional mortality of reef-building corals: the local-scale manifestations of global climate change and other local causes (McClanahan 2002, Hughes et al. 2003, Buddemeier et al. 2004, Aronson and Precht 2006, 2016, Rogers 2008, Côté and Darling 2010, Hoegh-Guldberg and Bruno 2010, Selig et al. 2012, Birkeland et al. 2013, Bruno and Valdivia 2016, Bruno et al. 2019). If the side-in impacts of diseases, hurricanes and bleaching continue unabated (Veron et al. 2009, Eakin et al. 2010, Hoegh-Guldberg et al. 2011, Randall et al. 2014, Manzello 2015, Precht et al. 2016), regional-scale coral mortality will outpace local recovery, resulting in reefs that will look and function very differently from the way they do now (Carpenter et al. 2008, Burman et al. 2012, Smith et al. 2013, Kuffner et al. 2014, Aronson and Precht 2016, Kuffner and Toth 2016, Precht et al. 2016, Toth et al. 2019). Macroalgae-dominated reefs could then truly become the norm.

## Acknowledgements

We thank the many people who contributed unpublished data or manuscripts in press. Portions of this work were carried out under permits from the Florida Keys National Marine Sanctuary and the Belize Department of Fisheries. Thanks to John Bruno, Steven Miller, and Lauren Toth for commenting on an earlier version of this manuscript. This research was supported by the UK Natural Environment Research Council (TAG); the Tyndall Centre for Climate Change Research (IMC, JAG, ARW); the US National Science Foundation (OCE-1535007 to RBA); the National Geographic Society; NOAA’s National Undersea Research Program, NOAA’s Sanctuaries and Reserves Division, Coastal Ocean Program, and FKNMS (RBA and WFP); and the Natural Science and Engineering Research Council of Canada (IMC). This is contribution number 146 from the Institute for Research to Global Climate Change of the Florida Institute of Technology.

