## Supplemental File 1 for "NON-RANDOM TIMING OF ECOLOGICAL SHIFTS ON CARIBBEAN CORAL REEFS SUGGESTS REGIONAL CAUSES OF CHANGE"

**Supplementary information**

**William F. Precht, Richard B. Aronson, Toby A. Gardner, Jennifer A. Gill, Julie P. Hawkins, Edwin A. Hernandez-Delgado, Walt C. Jaap, Tim R. McClanahan, Melanie D. McField, Thad J.T. Murdoch, Maggy M. Nugues, Callum M. Roberts, Christiane K. Schelten, Andrew R. Watkinson, and Isabelle M. Côté**

**Table S1**: See attached excel spreadsheet

**Supplementary Online Material – References associated with Table S1.**

Reference numbers refer to Study Code column in Table S1.

1. Porter, J. W., V. Kosmynin, K. L. Patterson, K. G. Porter, W. C. Jaap, J. Wheaton, K. Hackett, M. Lybolt, C. Tsokos, G. Yanev, D. M. Marcinek, J. Dotten, D. Eaken, M. E. Patterson, O. W. Meier, M. Brill, and P. Dustan. 2002. Detection of coral reef change by the Florida Keys Coral Reef Monitoring Project.pp 749-769 in J. W. Porter, and K. G. Porter, editors. The Everglades, Florida Bay, and Coral Reefs of the Florida Keys. An ecosystem sourcebook. CRC Press, Boca Raton.

Keller, B. D., editor. 2001. Sanctuary Monitoring Report 2000. U.S. Environmental Protection Agency, Florida.

Wheaton, J., W. C. Jaap, J. W. Porter, V. Kosminyn, K. Hackett, M. Lybolt, M. K. Callaham, J. Kidney, S. Kupfner, C. Tsokos, and G. Yanev. 2001. EPA/FKNMS coral reef monitoring project. Executive summary. Florida Fish and Wildlife Conservation Commission and University of Georgia, Athens.

2. Rogers, C. S., L. N. McLain, and C. R. Tobias. 1991. Effects of Hurricane Hugo (1989) on a coral-reef in St-John, USVI. *Marine Ecology-Progress Series* 78:189–199.

Rogers, C. S., V. Garrison, and R. Grober-Dunsmore. 1997. A fishy story about hurricanes and herbivory: seven years of research on a reef in St John, U.S. Virgin Islands. *Proceedings Eighth International Coral Reef Symposium* 1:555–560.

3 Personal communication with C Rogers, D Catanzaro, J Miller.

4 Bythell, J. C., M. Bythell, and E. H. Gladfelter. 1993a. Initial results of a long-term coral-reef monitoring program - Impact of Hurricane Hugo at Buck Island Reef National Monument, St-Croix, United-States Virgin-Islands. *Journal Experimental Marine Biology and Ecology* 172:171–183.

Bythell, J. C., E. H. Gladfelter, and M. Bythell. 1993b. Chronic and catastrophic natural mortality of 3 common Caribbean reef corals. *Coral Reefs* 12:143–152.

Bythell, J. C., E. H. Gladfelter, and Z. M. Hillis. 1993c. Ecological studies of Buck Island Reef National Monument, St. Croix, U.S. Virgin Islands: Report on February 1993 fixed transect surveys and post-Hurricane Hugo recovery of reef and coral community composition. US Department of Interior, National Park Service.

Bythell, J. C. 1998. Assessment of the impacts of hurricanes Marilyn and Luis and post-hurricane recovery at Buck Island Reef National Monument as part of the long-term coral reef monitoring program in the north-eastern Caribbean. US Department of Interior, National Park Service.

Bythell, J. C., Z. M. Hillis, B. Philips, W. J. Burnett, J. Larcombe, and M. Bythell. 2000a. Buck Island Reef National Monument, St Croix, US Virgin Islands: Assessment of the impacts of Hurricane Lenny (1999) and status of the reef in 2000. US Department of Interior, National Park Service.

Bythell, J. C., Z. M. Hillis-Starr, and C. S. Rogers. 2000b. Local variability but landscape stability in coral reef communities following repeated hurricane impacts. *Marine Ecology-Progress Series* 204:93–100.

5. Edmunds, P. J. 2002. Long-term dynamics of coral reefs in St John. *Coral Reefs* 21:357-367.

Edmunds, P. J., and J. D. Witman. 1991. Effect of Hurricane Hugo on the primary framework of a reef along the south shore of St-John, United-States Virgin-Islands. *Marine Ecology-Progress Series* 78:201–204.

6. Edmunds, P. J. 2002. Long-term dynamics of coral reefs in St John. *Coral Reefs* 21:357–367.

7. Nemeth, R. S. 1999. Interim report on the effects of the Caret Bay Villas Construction on the coral reef environment at Caet Bay, St Thomas, USVI. University Virgin Islands, St Thomas.

Nemeth, R. S. 2001. The effect of sedimentation from coastal development on coral bleaching. University Virgin Islands, St Thomas, USVI.

Nemeth, R. S., and J. S. Nowlis. 2001. Monitoring the effects of land development on the near-shore reef environment of St. Thomas, USVI. *Bulletin Marine Science* 69:759–775.

8. Liddell, W. D., and S. L. Ohlhorst. 1986. Changes in benthic community composition following the mass mortality of *Diadema* at Jamaica. *Journal Experimental Marine Biology Ecology* 95:271–278.

Liddell, W. D., and S. L. Oldhorst. 1992. Ten years of disturbance and change on a Jamaican fringing reef. *Proceedings Seventh International Coral Reef Symposium*:144–150.

9. Hughes, T. P. 1989. Community structure and diversity of coral reefs - the role of history. *Ecology* 70:275–279.

Hughes, T. P. 1994. Catastrophes, phase-shifts, and large-scale degradation of a Caribbean coral-reef. *Science* 265:1547–1551.

Hughes, T. P. 1996. Demographic approaches to community dynamics: A coral reef example. *Ecology* 77:2256–2260.

Hughes, T. P., and J. H. Connell. 1999. Multiple stressors on coral reefs: A long-term perspective. *Limnology Oceanography* 44:932–940.

Hughes, T. P., and J. B. C. Jackson. 1985. Population-dynamics and life histories of foliaceous corals. *Ecological Monographs* 55:141–166.

10. Aronson, R. B., and W. F. Precht. 2000. Herbivory and algal dynamics on the coral reef at Discovery Bay, Jamaica. *Limnology Oceanography* 45:251–255.

11. Aronson, R. B., and W. Precht, F. 2001. Evolutionary paleoecology of Caribbean coral reefs. Pages 171-233 in W. D. Allmon, and D. J. Bottjer, editors. Evolutionary paleoecology: the ecological context of macroevolutionary change. Columbia University Press, New York.

12. Knowlton, N., J. C. Lang, M. C. Rooney, and P. A. Clifford. 1981. Evidence for delayed mortality in hurricane-damaged Jamaican staghorn corals. *Nature* 294:251–252.

Knowlton, N., J. C. Lang, and B. D. Keller. 1990. Case study of natural population collapse: post hurricane predation on Jamaican staghorn corals. *Smithsonian Contributions to Marine Science* 31:1–25.

13. Edmunds, P. J., and J. F. Bruno. 1996. The importance of sampling scale in ecology: kilometer-wide variation in coral reef communities. *Marine Ecology-Progress Series* 143:165–171.

Cho, L. L., and J. D. Woodley (2002). Recovery of reefs at Discovery Bay, Jamaica and the role of *Diadema antillarium*. *Proceedings Ninth International Coral Reef Symposium:* 331-338.

14. Andres, N. G., and J. D. Witman. 1995. Trends in community structure on a Jamaican reef. *Marine Ecology-Progress Series* 118:305-310.

Cho, L. L., and J. D. Woodley (2002). Recovery of reefs at Discovery Bay, Jamaica and the role of *Diadema antillarium*. *Proceedings Ninth International Coral Reef Symposium*: 331–338.

15. Gayle, P. M. H., and J. D. Woodley. 1998. Discovery Bay, Jamaica. Pp. 17–33 in B. Kjerfve, editor. CARICOMP - Caribbean coral reef, seagrass and mangrove sites. UNESCO, Paris.

16. McClanahan, T. M., R. B. Aronson, W. Precht, F., and N. A. Muthiga. 1999. Fleshy algae dominate remote coral reefs of Belize. *Coral Reefs* 18:61–62.

McClanahan, T. R., and N. A. Muthiga. 1998. An ecological shift in a remote coral atoll of Belize over 25 years. *Environmental Conservation* 25:122–130.

17. McClanahan, T. M. 2001. Glover's Reef: Patch Reef Monitoring Program - Status in 2001. Wildlife Conservation Society.

McClanahan, T. R., M. McField, M. Huitric, K. Bergman, E. Sala, M. Nystrom, I. Nordemar, T. Elfwing, and N. A. Muthiga. 2001. Responses of algae, corals and fish to the reduction of macroalgae in fished and unfished patch reefs of Glovers Reef Atoll, Belize. *Coral Reefs* 19:367–379.

18. McField, M. 2001. The influence of disturbance and management on coral reef community structure in Belize. PhD Thesis. University of South Florida, St Petersburg.

19. Aronson, R. B., and W. Precht, F. 2001. Evolutionary paleoecology of Caribbean coral reefs. Pages 171–233 in W. D. Allmon, and D. J. Bottjer, editors. Evolutionary paleoecology: the ecological context of macroevolutionary change. Columbia University Press, New York.

Aronson, R. B., W. Precht, F., M. A. Toscano, and K. H. Koltes. 2002. The 1998 bleaching event and its aftermath on a coral reef in Belize. *Marine Biology* 141:435–447.

20. Koltes, K. H., J. Tschirky, and I. C. Feller. 1998. Carrie Bow Caye, Belize. pp 79-94 in B. Kjerfve, editor. CARICOMP - Caribbean coral reef, seagrass and mangrove sites. UNESCO, Paris.

Rutzler, K., and I. G. Macintyre. 1982. The habitat distribution and community structure of the Barrier Reef complex at Carrie Bow Cay, Belize. pp 9–45 in K. Rutzler, and I. G. Macintyre, editors. The Atlantic barrier reef ecosystem at Carrie Bow Cay, Belize, I. Structure and communities. Smithsonian Institution Press, Washington.

21. Garrison, V., E. A. Shinn, J. Miller, M. Carlo, R. Rodriguez, and K. H. Koltes. 2000. Isla de Culebra, Puerto Rico changes in benthic cover on three reefs (1991–1998). Technical report for the Water Resources Division of the U.S. Geological Survey, St Petersburg.

Shinn, E. A., and R. B. Halley. 1992. Assessment of damage to coral reefs by Hurricane Hugo in W. C. Schwab, and R. W. Rodriguez, editors. Progress of studies on the impact of Hurricane Hugo on the coastal resources of Puerto Rico. United States Geological Survey Open-File Report OF 92-0717.

22. Hernandez-Delgado, E. A. 2000. Effects of anthropogenic stress gradients in the structure of coral reef fish and epibenthic communities. Faculty of Natural Sciences. University Puerto Rico, Rio Pedras Campus, San Juan.

Hernandez-Delgado, E. A. 2001. Effects of the Luis Pena Channel Marine Fishery Reserve, (Culebra Island) in the structure of coral reef epibenthic communities: I Ecological change of coral reefs (1997–2001). Technical Report, Puerto Ricao Coastal Zone Management Program - Department of Natural and Environmental Resources, San Juan, Puerto Rico.

23. Garcia, J. R., C. Schmitt, C. Heberer, and A. Winter. 1998. La Parguera, Puerto Rico, USA. Pages 195–212 in B. Kjerfve, editor. CARICOMP - Caribbean coral reef, seagrass and mangrove sites. UNESCO, Paris.

24. Ruiz-Renteria, F., B. I. van Tussenbroek, and E. Jordan-Dahlgren. 1998. Puerto Morelos, Quintana Roo, Mexico. Pages 57–66 in B. Kjerfve, editor. CARICOMP - Caribbean coral reef, seagrass and mangrove sites. UNESCO, Paris.

25. Rodriguez, R. W., and E. Jordan-Dahlgren. 1996. Short-term effects of Hurricane Roxanne on a littoral coral community in Cozumel, Mexico. Unpublished manuscript.

26. Garza-Perez, J. R., and J. E. Arias Gonzalez. 2001. Temporal changes of a coral reef community in the South Mexican Caribbean. Pages 415–427 in R. LeRoy Creswell, editor. Proceedings of the 52nd Annual Gulf and Caribbean Fisheries Institute., Key West, Florida Nov. 1999.

27. De Meyer, K. 1998. Bonaire, Netherland Antilles. Pages 141–149 in B. Kjerfve, editor. CARICOMP - Caribbean coral reef, seagrass and mangrove sites. UNESCO, Paris.

28. Personal communication with C Glendinning.

29. Gerace, D., G. K. Ostrander, and G. W. Smith. 1998. San Salvador, Bahamas. Pages 220–245 in B. Kjerfve, editor. CARICOMP - Caribbean coral reef, seagrass and mangrove sites. UNESCO, Paris.

Ostrander, G. K., K. M. Armstrong, E. T. Knobbe, D. Gerace, and E. P. Scully. 2000. Rapid transition in the structure of a coral reef community: The effects of coral bleaching and physical disturbance. *Proceedings National Academy Sciences USA* 97:5297–5302.

45. Personal communication with C. Schelten.
